# Ribosome profiling to study translation in cortical synaptoneurosomes on group 1 mGluR activation

**DOI:** 10.64898/2026.05.10.723250

**Authors:** Preeti Madhav Kute, Kornel Labun, Håkon Tjeldnes, Eivind Valen, Ravi Muddashetty

## Abstract

Local protein synthesis in neurons occurs in both axons and dendrites and plays a central role in synaptic function. High-throughput-based sequencing and imaging studies have demonstrated the presence and translation of synaptically localised mRNAs. However, quantification of activity-dependent translation dynamics at synapses at the transcriptome-wide scale remains limited. Here, we apply ribosome profiling to synapse-enriched fractions (synaptoneurosomes) derived from rat cortical tissue following stimulation with the group 1 mGluR agonist DHPG. DHPG stimulation induced translation of mRNAs involved in synaptic processes, including synaptic vesicle exocytosis and axo-dendritic transport. Notably, translation of ribosomal protein mRNAs was upregulated upon mGluR activation, consistent with the expected increase in *de novo* protein synthesis. Together, these results demonstrate the use of ribosome profiling to capture changes in local mRNA translation from isolated preparations.

## Introduction

Activation of group 1 metabotropic glutamate receptors (mGluR1 and mGluR5) with 3,5-dihydroxyphenylglycine (DHPG) is known to increase *de novo* protein synthesis at neuronal synapses (Muddashetty et al. 2007; Raymond et al. 2000; Ghosh Dastidar et al. 2020; Mockett et al. 2011; Mango & Ledonne 2023). This *de novo* protein synthesis is required for the induction and maintenance of a form of synaptic plasticity called mGluR-dependent long-term depression (mGluR-LTD) in the CA1 region of the mouse hippocampus, and involves key transcripts such as *Arc, Fmr1*, and MAP1b (Huber et al. 2000; Waung et al. 2008; Park et al. 2008; Weiler et al. 1997; Hou et al. 2006). Further, this response is mediated by the activation of the ERK and mTOR pathways, which promote translation initiation and elongation (Hou & Klann 2004; Gallagher et al. 2004; Sharma et al. 2010). These effects are mediated through increased formation of eIF4F complex, elevated ribosomal protein S6 phosphorylation, and reduced phosphorylation of eEF2 (Banko et al. 2006; Antion et al. 2008; Ghosh Dastidar et al. 2020) but also increased eIF2α phosphorylation (Di Prisco et al. 2014).

Several studies have sought to identify the translatome underlying mGluR-mediated synaptic plasticity using different high-throughput approaches. Polysome profiling coupled with RNA-seq in cortical neuronal cultures identified translationally upregulated mRNAs following DHPG treatment, including *Ophn1* that was found to be required for mGluR-LTD through its role in AMPAR surface expression (Di Prisco et al. 2014). Complementary approaches combining Translating Ribosome Affinity Purification (TRAP-seq) and RNA-seq, Seo et al. revealed translation upregulation of ribosomal protein and mitochondrial mRNAs upon induction of mGluR-LTD in the CA1 region of the mouse hippocampus (Seo et al. 2022). Ribosome profiling in DHPG-treated mouse hippocampal slices further identified upregulation of transcripts related to GPCR activity, synaptic membrane, and endoplasmic reticulum membrane (Hien et al. 2020). At the proteome level, quantitative peptidomics using AHA labelling and tandem mass tagging demonstrated increased abundance of ribosomal and translation-related proteins upon mGluR activation (van Gelder et al. 2020).

Despite these advances, most studies address the translation response in bulk samples using either hippocampal cultures or slices, whereas mGluR-activation response initiates locally at the synaptic sites (Hafner et al. 2019; Sun et al. 2021). Translation of synaptically-localised mRNAs are induced within minutes of receptor activation, contributing to the local proteome (Cajigas et al. 2012; Glock et al. 2021; Donlin-Asp et al. 2021). Analyses at the tissue- or culture-level may therefore obscure the role of spatially restricted, activity-mediated translation at synaptic sites.

Synaptoneurosomes, a simple and rapid isolation of pre- and post-synaptic terminals that retain neurochemical properties, provide a tractable system to address this limitation, enabling study of local translation decoupled from transcription and RNA transport (Chang et al. 2012; Simbriger et al. 2020). Earlier transcriptomics and translatomics analyses, using naïve, untreated synaptic preparations to characterise synaptic mRNAs, have shown the presence of ribosome-bound mRNAs in such a system (Williams et al. 2009; Simbriger et al. 2020; Ouwenga et al. 2017) and protein synthesis following neuroreceptor activation (Kuzniewska et al. 2020; Chmielewska et al. 2019; Muddashetty et al. 2007; Scheetz et al. 2000; Yin et al. 2002; Kute et al. 2019). However, an unbiased transcriptome-wide assessment of receptor activity-dependent translation is lacking, likely due to the heterogeneous nature and the low abundance of RNA. Such an approach, ribosome profiling, was applied to untreated synaptoneurosomes, showing enrichment of synaptically relevant transcripts (Simbriger et al. 2020) upon synaptic stimulation.

Here, we extend previous work by combining synaptoneurosome isolation and ribosome profiling to directly detect activity-mediated translation. We generated high-quality ribosome profiling libraries that showed the characteristics of 28-nt ribosome footprints, triple nucleotide periodicity, and mapping to the coding region. Differential expression analysis revealed translation upregulation of ribosomal protein mRNAs and transcripts associated with synaptic function upon mGluR activation.

This proof-of-concept study establishes ribosome profiling of stimulated synaptoneurosomes as a framework for characterizing the regulation of local translation at synapses.

## Results

### Ribosome profiling in synaptoneurosomes post group 1 mGluR activation

Previous studies have shown that stimulation of synaptoneurosomes with DHPG, an agonist of group 1 mGluRs, leads to a decrease in eEF2 phosphorylation levels and a concomitant increase in protein synthesis, consistent with enhanced translation elongation (Ghosh Dastidar et al. 2020; Muddashetty et al. 2007; van Gelder et al. 2020). To test the effect of group 1 mGluR activation at the transcriptomic level, we performed ribosome profiling (Ingolia et al. 2012) on synaptoneurosomes treated without (basal) or with DHPG (50 μM, 5 min, 37°C). The synaptoneurosomes have been previously validated for their enrichment of post-synaptic protein PSD-95 and their snowman-shaped structure (**Figure 1a**, EM image) (Ghosh Dastidar et al. 2020; Kute et al. 2019). Poly-A selected RNA-seq libraries (unmatched) were also prepared to account for changes in RNA abundance.

**Figure 1:**
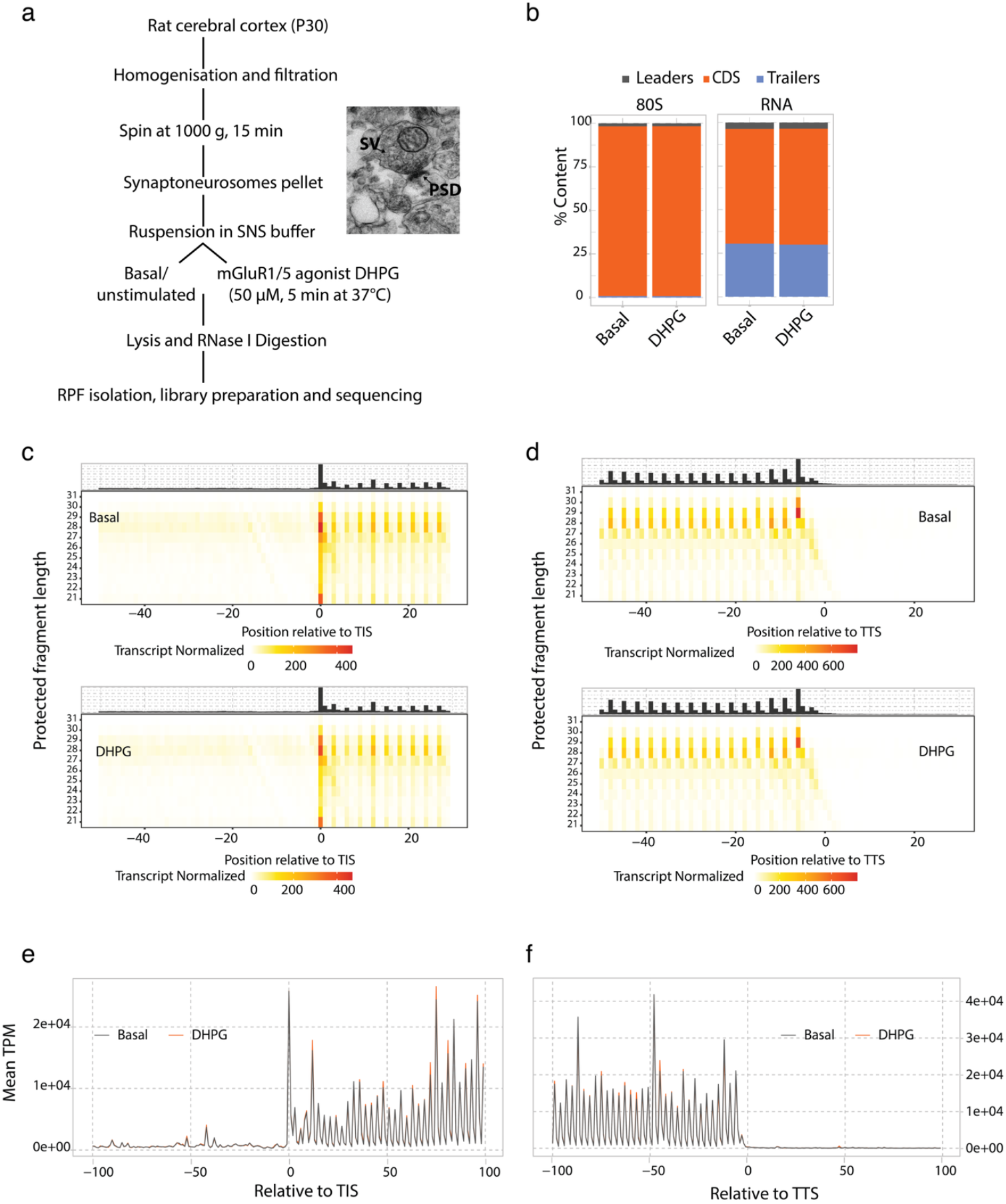
Ribosome profiling from rat cortical synaptoneurosomes after synaptic stimulation with DHPG. a. Schematic describing the steps to obtain ribosome protected fragments (RPFs) from rat synaptoneurosomes after mGluR activation, with a representative EM image of synaptoneurosome (PSD-postsynaptic density, SV-synaptic vesicles) b. Percent mapping of transcriptomic reads to the regions of the mRNA: leaders (5’UTR), CDS, and trailers (3’ UTR) for ribo-seq and RNA libraries. c. RPFs around translation initiation sites (TIS) from basal and DHPG-treated samples d. RPFs around translation termination sites (TTS) from basal and DHPG-treated samples e. Read distribution for region 100 nt upstream and downstream of the TIS for both basal and DHPG-treated samples. TPM: transcript per million f. Read distribution for region 100 nt upstream and downstream of the TTS for both basal and DHPG-treated samples. TPM: transcript per million

For both modalities, we observed a high level of reproducibility within basal and DHPG-treated conditions (**Supplementary Figure 1a-1d**). Ribosome-protected fragments (RPF) displayed the expected read length distribution from 26-29, peaking at 28 nt (**Supplementary Figure 1e**). More than 75% of RPF reads aligned to the mRNA, compared to ∼40% for the RNA-seq reads (**Supplementary Figure 1f**). Mapping across transcript regions showed that more than 90% of RPFs localized to coding regions (CDS), with the RNA-seq more broadly distributed, with 50% to the CDS and around 25% to the trailers (**Figure 1b**). RPF reads of both basal and DHPG-treated synaptoneurosomes showed the expected 3-nt periodicity, which was absent for the RNA-seq reads (**Supplementary Figure 1g**). The periodicity was preserved across footprints lengths (21-31 nt) and observed over the translation initiation site (TIS, **Figure 1c**) and termination site (TTS, **Figure 1d**) Metagene analysis further demonstrated a relatively uniform distribution of ribosomes throughout the CDS (**Figure 1e and 1f**). Together, these results demonstrate the successful capture of high-quality ribosome footprints from rat brain synaptoneurosomes.

### mGluR activation differentially affects synaptic translation

To explore the effect of mGluR activation on translation of individual mRNAs, we used DESeq2 to identify differentially expressed transcripts (Love et al. 2014) in the riboseq libraries. This identified 80 transcripts that were significantly differentially regulated upon DHPG-treatment compared to the basal state (adjusted p < 0.05). Of these, 26 were upregulated, and 54 were downregulated (**Figure 2a, Supplementary Table 1**). Among the most strongly upregulated transcripts were *Csnk1d, Rpl38-ps8, Mcfd2, Ptma*, and *Ptmal5*, while the top downregulated candidates were *Mmd, Dpp10, Fads1, Acad8*, and *Cers5*. There were no global differences in translation efficiency (**Supplementary Figure 2a**) and RNA abundance (**Supplementary Figure 2b**) between the basal and DHPG conditions.

**Figure 2:**
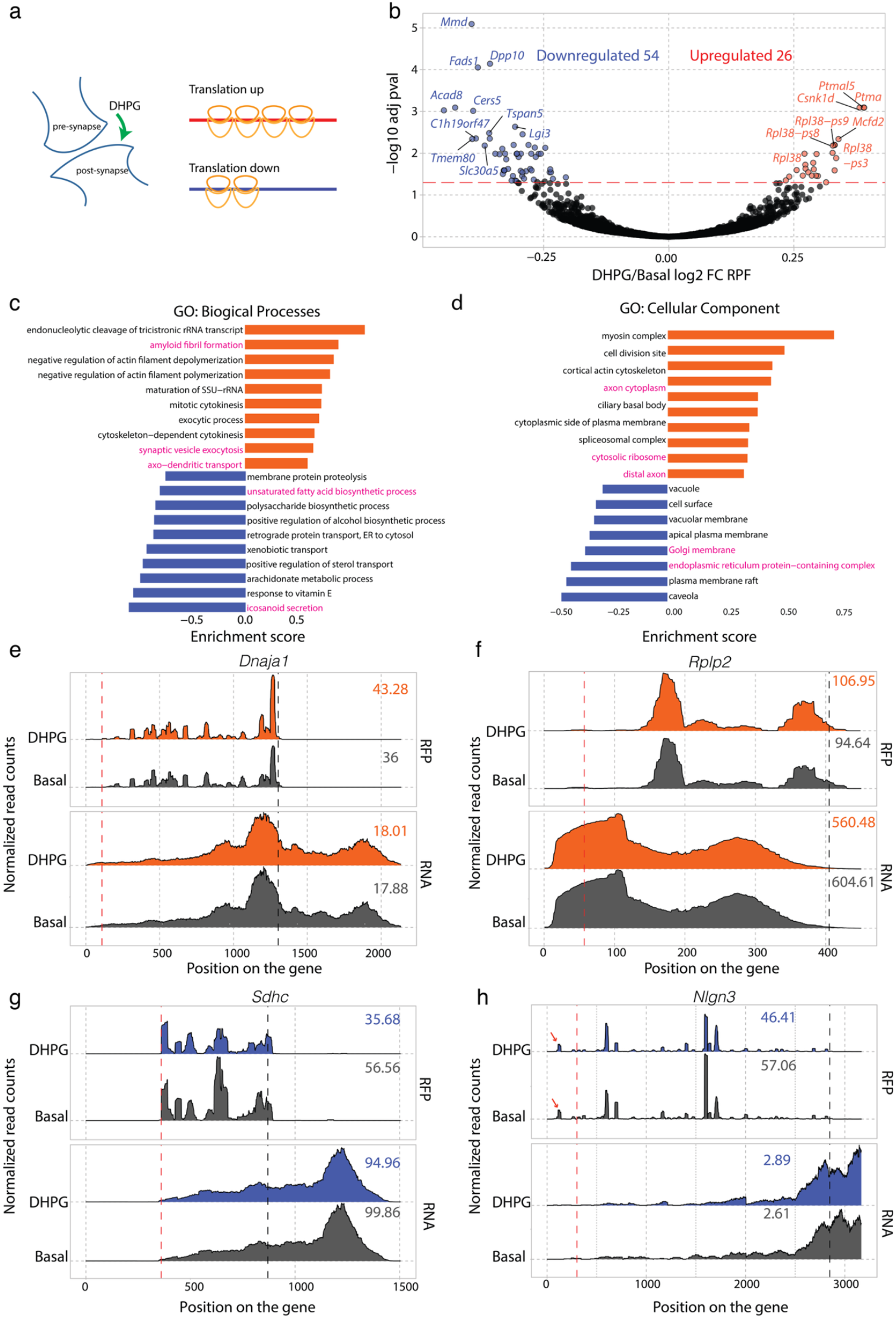
Differential translation of genes on DHPG stimulation in synaptoneurosomes. a. Schematic showing DHPG-mediated translation dynamics at synapses. b. RPF-based fold change between DHPG and basal, labelled with top differentially expressed transcripts (adjusted p-value < 0.05) with no fold change cut-off. c. Gene ontology analysis for differentially expressed transcripts in the biological process category. Red bars are upregulated genes, and blue bars are downregulated genes. Highlighted categories are mentioned in the text d. Gene ontology analysis for differentially expressed transcripts in the Cellular components category. Red bars are upregulated genes, and blue bars are downregulated genes. highlighted categories are mentioned in the text e. Gene silhouettes for DnaJ Heat Shock Protein Family (Hsp40) Member A1 (*Dnaja1*), upregulated on DHPG stimulation, for RPFs and RNA-seq libraries. Red and black dotted lines indicate translation start and termination sites, respectively. f. Gene silhouettes for Ribosomal Protein Lateral Stalk Subunit P2 (*Rplp2)*, upregulated on DHPG stimulation, for RPFs and RNA-seq libraries. Red and black dotted lines indicate translation start and termination sites, respectively. g. Gene silhouettes for succinate dehydrogenase complex subunit C (*Sdhc*), downregulated on DHPG stimulation, for RPFs and RNA-seq libraries. Red and black dotted lines indicate translation start and termination sites, respectively. h. Gene silhouettes for Neuroligin3 (*Nlgn3*), downregulated on DHPG stimulation, for RPFs and RNA-seq libraries. Red and black dotted lines indicate translation start and termination sites, respectively. Arrows indicate an active uORF.

Gene ontology analysis revealed enrichment of both pre- and post-synaptic processes following DHPG stimulation, including synaptic vesicle exocytosis, axo-dendritic transport, and amyloid fibril formation, whereas processes such as eicosanoid secretion and retrograde protein transport were depleted (**Figure 2b**). Consistent with this, cellular component analysis showed enrichment of synaptic compartments, including axonal cytoplasm and the cytosolic ribosome, alongside depletion of endoplasmic reticulum- and Golgi-associated categories. Protein-protein interaction analysis of the 80 differentially regulated transcripts using the STRING database in Cytoscape (Shannon et al. 2003) revealed a prominent cluster of ribosomal proteins (**Supplementary figure 2c**). Among these mRNAs such as *Rpl38, Rpl23*, and *Rplp2*, were upregulated following DHPG treatment, whereas mitochondrial transcripts such as *Cox6b1, Sdhc*, and ND5 were downregulated (**Supplementary figure 2c**).

Inspection of individual transcripts supported these global trends. Representative upregulated gene transcripts, DnaJ Heat Shock Protein Family (Hsp40) Member A1 (*Dnaja1*) and Ribosomal Protein Lateral Stalk Subunit P2 (*Rplp2)*, showed an increase in RPF counts on DHPG stimulation without corresponding differences in RNA abundance (**Figure 2e** and **2f**). In contrast, downregulated gene candidates succinate dehydrogenase complex subunit C (*Sdhc*) and Neuroligin3 (*Nlgn3*) show a reduction in RPF counts upon DHPG-treatment (**Figure 2f** and **2g**).

Given the heterogeneous preparation of synaptic, nonsynaptic, and non-neuronal structures, glial contamination is expected. Accordingly, we detect glial genes such as *Gfap, Slc1a2* and *Aqp4* for both RFP and RNA-seq libraries (**Supplementary Figures 2d-2f**). However, the availability of datasets for neuronal and glial-specific genes (Glock et al. 2021; Kaulich et al. 2025) enables filtering of these genes to focus on the translation of neuronally- and synaptically enriched genes. Together, these results demonstrate that ribosome profiling of synaptoneurosomes can robustly capture transcript-specific changes in translation.

## Discussion

In this study, we employed a rapid and reproducible method to isolate pre- and post-synaptic structures called synaptoneurosomes from the adult rat brain cortex (Muddashetty et al. 2007). The preparations retain metabolic, enzymatic, and biochemical function and activity (Ghosh Dastidar et al. 2020; Muddashetty et al. 2011; Kuzniewska et al. 2018; Kute et al. 2019). Using this approach, we demonstrate the presence and active translation of synaptically relevant mRNAs on group 1 mGluR activation. Analysis of RPFs revealed both a decrease and an increase in ribosomal occupancy on mRNAs, implying bidirectional translation regulation, as has also been observed in other high-throughput studies on DHPG-mediated translation response (Hien et al. 2020; Seo et al. 2022).

mGluR activation is known to activate the Akt-mTOR pathway (Sharma et al. 2010; Hou & Klann 2004) and the synthesis of 5′ oligopyrimidine tract (5′ TOP) protein EF1A (Antion et al. 2008). While we did not check the activation of mTOR in this study, the increase in ribosomal occupancy of some of the ribosomal mRNAs (*Rpl38, Rpl23, Rplp2*) which contain 5′ TOP sequence is consistent with mTOR activation in synaptoneurosomes on DHPG stimulation. This increase in translation of ribosomal mRNAs also aligns with other translatomic studies in mouse hippocampus (Seo et al. 2022; van Gelder et al. 2020).

Recent studies have shown rapid translation and incorporation of ribosomal proteins into existing ribosomes at the neurites (Fusco et al. 2021; Fusco et al. 2025), independent of the canonical ribosome biogenesis occurring in the nucleus. Our observation of rapid translation of neurite-localised ribosomal mRNAs upon mGluR activation supports the hypothesis of incorporation of nascent ribosomal proteins into existing ribosomes to maintain protein synthesis required for mGluR-LTD (Seo et al. 2022). In contrast to Seo et al., who observed an upregulation of mitochondrial mRNAs, in our study, we found downregulation of some of the mitochondrial mRNAs (e.g. *Cox6b1*, ND5, and *Sdhc*). This discrepancy could be explained by differences in methods and experimental design. While Seo et al. perform TRAP-seq with a 25 min washout period, we immediately lyse and digest the lysates after DHPG stimulation. Potentially, we are detecting early translation of some of the transcripts, such as ribosomal protein mRNAs, through redistribution of ribosomes from other mRNAs, such as mitochondrial mRNAs.

### Limitations and future perspective

The conditions for the DHPG treatment may not be optimal (50 μM, 5 min at 37°C). Increasing both time and dose could produce a stronger and more robust response, as demonstrated previously (van Gelder et al. 2020; Hien et al. 2020). Similarly, synaptoneurosomes can be incubated longer at 37°C after stimulation and before lysis to obtain a more robust translation response (Ghosh Dastidar et al. 2020; Seo et al. 2022). Some studies have incubated the untreated synaptoneurosomes at 4°C, while incubation of the treated group was at 37°C (Kuzniewska et al. 2018), potentially producing a different translation response relative to untreated samples. In addition, the lack of aeration could induce metabolic stress, potentially contributing to the reduced translation of mitochondrial mRNAs.

Another limitation is the lack of matched RNA-seq libraries along with ribosome profiling, based on the assumption that stimulation would not affect the transcriptome of synaptoneurosomes. While this is likely valid for a rapid response, it limits the exact quantification of translational efficiency.

While synaptoneurosomes are rich in synaptic content and protein synthesis machinery to study rapid translation and post-translation response, such as phosphorylation (occurring within minutes), they are not useful for long-term processes, like the role of translation on synapse structure, neurite growth, where neuronal cultures turn out to be a better system.

Despite these limitations, synaptic preparations are a powerful system to study synaptic plasticity mechanisms, especially in physiologically relevant disease contexts in both animals and humans (Ahmad & Liu 2020). Notably, synaptoneurosomes can be prepared from frozen tissues such as human brain samples (Franklin & Taglialatela 2016) opening opportunities to study translation regulation in clinical samples using ribosome profiling. In addition, emerging techniques such as ribosome complex profiling (RCP-seq) and selective translation complex profiling (sel-TCP-seq) enable the characterization of translation initiation dynamics (Giess et al. 2020; Kute et al. 2025; Wagner et al. 2022). Applying these to synaptoneurosomes can shed light on the role of initiation factors, such as eIF4E and eIF2ɑ, in regulating synaptic translation.

## Materials and Methods

### Animal ethics

All the work was done with due approval from the Institutional Animal Ethics committee (IAEC) and the Institutional BioSafety Committee (IBSC), InStem, Bangalore, India

### Synaptoneurosome preparation

Sprague Dawley (SD) male rats from the age between post-natal day 28–33 (P28-33) were used for synaptoneurosome preparation. Rat cerebral cortex was dissected, with the olfactory lobe and the striatum removed from the cortex. The tissue was homogenized using a dounce homogenizer in ice in 10 volumes (∼10ml) of synaptoneurosome buffer (25 mM Tris–HCl pH 7.4, 118 mM NaCl, 4.7 mM KCl, 1.2 mM MgSO_4_, 2.5 mM CaCl_2_, 1.53 mM KH_2_PO_4_, and 212.7 mM glucose, supplemented with protease inhibitors (Roche), and RNaseOut (30 U/mL). Homogenates were then passed through three 100-μm nylon mesh filters followed by one 11-μm filter MLCWP 047 Millipore (Bedford, MA). The filtrate was centrifuged at 1,000 *g* for 15 min at 4°C. The pellet was resuspended in 2 mL of the synaptoneurosome buffer.

### Synaptoneurosome stimulation and lysis

For stimulation, synaptoneurosomes were pre-warmed at 37°C for a minimum of 5 min at 1300 rpm in a thermal block. Synaptoneurosomes were split into two of 1 mL each. 50 μM DHPG (made in nuclease free water) was added to one sample, and the other sample was left untreated and both were incubated for 5 min at 37°C at 1300 rpm. Stimulation was stopped by pelleting the synaptoneurosomes at 5000 rpm for 1 min and resuspending in 1 mL lysis buffer (50 mM Tris pH 7.5, 150 mM NaCl, 5 mM MgCl_2_, 1% NP-40, 0.1 mg/mL cycloheximide, protease inhibitor cocktail). The lysate was clarified by centrifugation at 20000 *g* for 20 min at 4 C and supernatant was collected.

### Ribosome profiling

Ribosome profiling was done from the synaptoneurosomal lysates according to Ingolia et. al. (Ingolia et al. 2012) with few modifications. Ribosome profiling was done in three replicates using three animals. After stimulation with DHPG and lysis, 800 µL of lysate (out of 1.0 mL) was digested with RNase I (100 U/µL; Invitrogen, AM2294) for 45 min at room temperature. The RNase-treated lysate was then layered on 1M sucrose solution and spun at 70,000 RPM for 4h in TLA 100.3 rotor (Beckman Coulter). The ribosome pellet obtained was then dissolved in TRIzol for RNA extraction. Extracted RNA was then run on 12% urea-PAGE for 2h at 200V along with oligo markers. Bands between 25-34 nt were cut, and RNA was eluted out from the gel using 500 µl of 0.3 M NaCl overnight. Precipitated RNA was then dephosphorylated at the 3’ end with PNK for 1h at 37°C and phosphorylated at the 5’ end with PNK and 1 mM ATP for 1 h at 37°C. Libraries were prepared using Illumina TruSeq Small RNA sample preparation kit. Libraries were size selected on acrylamide gel and quality checked on Tapestation for size and concentration. Libraries were sequenced on the Nextseq Illumina sequencer to obtain 120 million reads of 75 single end per library.

### RNA seq libraries

Unmatched RNA seq analysis was done from synaptoneurosome lysates post DHPG treatment (n=2 independent replicates). RNA was isolated from the lysates using the TRIzol method and resuspended in 10 ul of nuclease free water. Library preparation was done by the Poly-A enrichment method and sequenced on Illumina Hiseq 2500 with 100 bp paired end sequencing. Approximately 42 million reads were obtained per sample.

### Ribosome profiling analysis

The repository at Valen lab / Preeti_Jain_2020_Rattus_norvegicus · GitLab contains all the code that was used for analysis, and all the processed data and figures are available there for inspection. All figures and processed tables are present in the repository. For the data processing, we used ORFik (1.29.11) as shown in the script “0_preprocess.R”. Analysis uses the latest at the time, ensembl Rattus norvegicus (GRCr8 / GCA_036323735.1). ORFik pipeline uses fastp software for trimming and STAR for alignment. Paired-end fastq reads are initially aligned to the contaminants - phix, rRNA, ncRNA, tRNA, and finally to the genome. Important options that were set are trim.front = 0, min.length = 20. Aligned data is processed using ORFik and custom scripts available in the data repository. We restricted genes with multiple transcripts to a single transcript by selecting the one with the longest coding sequence. For statistical testing, we used 0.05 as the significance level (p adjusted).

For comparisons at the global-level, we calculate measures produced per library using FPKM values instead of raw counts and average replicates, furthermore, we normalise using reads per million reads for that group of genes to gain a comprehensive perspective. For Translation efficiency (TE) calculations we divided counts of RPFs on CDS by counts of RNA on mRNA.

### Statistics

Riboseq experiments were performed using three biological replicates, i.e., three different male rats. For RNAseq, only two replicates were conducted. For boxplot figures t-test statistics were used to compare means of the distributions with corrected p-values using Benjamini-Hochberg normalization.

## Data availability

Raw data from one replicate is hosted as BioProject under PRJNA386439. All three replicates are uploaded and processed on ribocrypt.org under the project ID PRJNA386439 and sd_preeti2.

## Author contributions

PMK and RSM designed and formulated the experiments. KL and HT conducted the bioinformatic analysis. PMK, KL and EV interpreted the data. PMK and EV wrote the manuscript with inputs from all the authors.

## Acknowledgements

We would like to thank the Sequencing facility and Animal House Facility at the National Centre for Biological Sciences (NCBS), Bangalore, India. We would also like to acknowledge Michał Świrski’s (University of Warsaw, Poland) help with uploading the data on ribocrypt.org.

## Conflict of Interest Statement

The authors declare that they have no conflict of interest.

## Notes

https://git.app.uib.no/valenlab/preeti_jain_2020_rattus_norvegicus

### Competing Interest Statement

The authors have declared no competing interest.

