## Supplementary data for "Ribosome profiling to study translation in cortical synaptoneurosomes on group 1 mGluR activation"

**Supplementary figures and table**

Supplementary figure 1 (related to figure 1): Quality check of riboseq libraries from rat synaptoneurosomes

Supplementary figure 2 (related to figure 2): Differential gene analysis at the TE and RNA level, cytoscape

Supplementary Table 1: List of differentially regulated transcripts at the RPF level

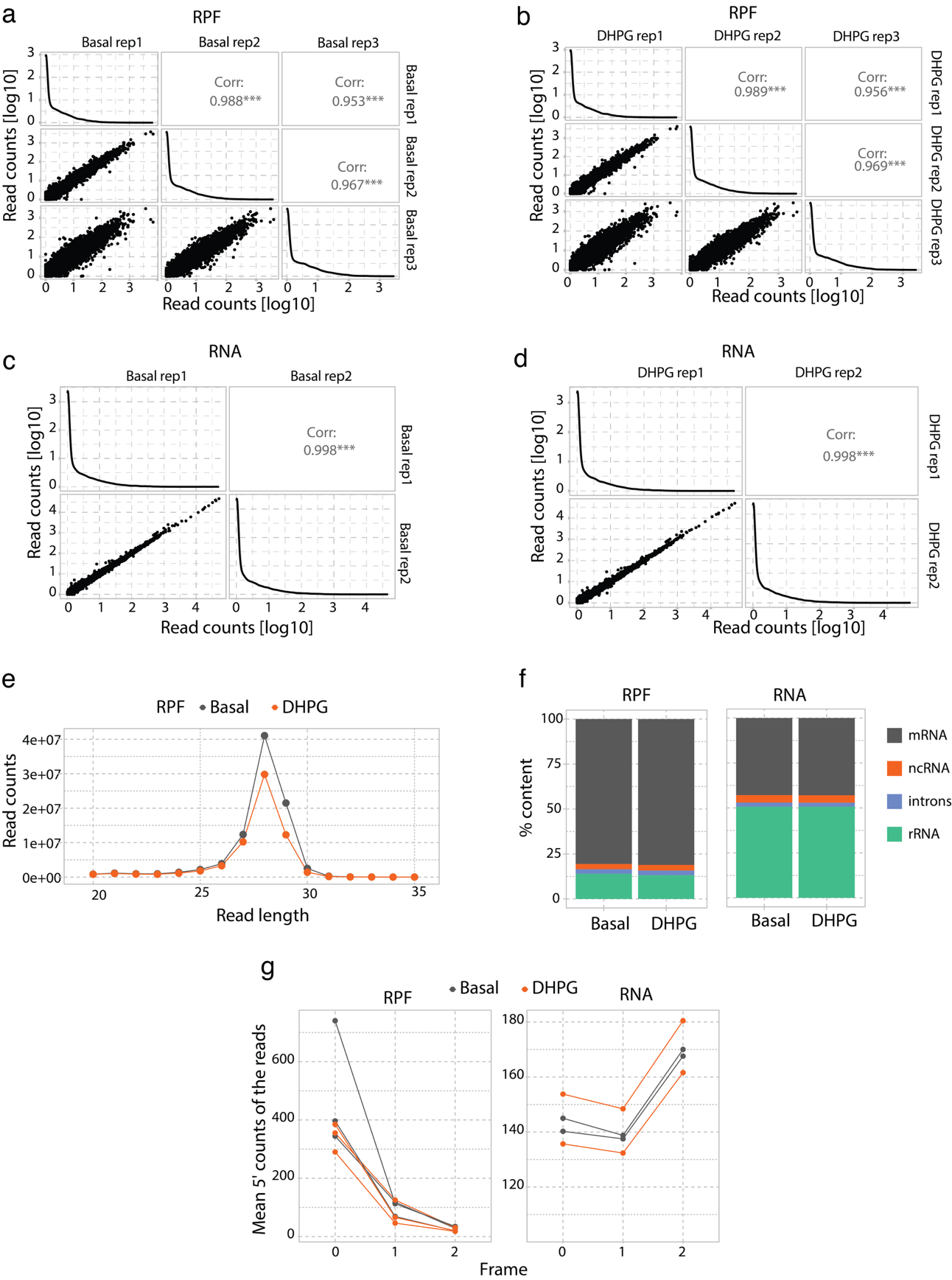

**Supplementary figure 1 (related to figure 1): Quality check of riboseq libraries from rat synaptoneurosomes**

a. Spearman’s correlation analysis of RPF libraries for basal samples.
a. Spearman’s correlation analysis of RPF libraries for DHPG treated samples.
a. Spearman’s correlation analysis of RNA libraries for basal samples.
a. Spearman’s correlation analysis of RNA libraries for DHPG treated samples.
a. RPF distribution for basal and DHPG treated samples
a. Percent mapping of sequenced reads to RNA biotypes such as mRNA, non-coding RNA (ncRNA), introns, and rRNA for basal and DHPG groups for RPF and RNA libraries.
a. Three-frame distribution of reads for RPF and RNA libraries.

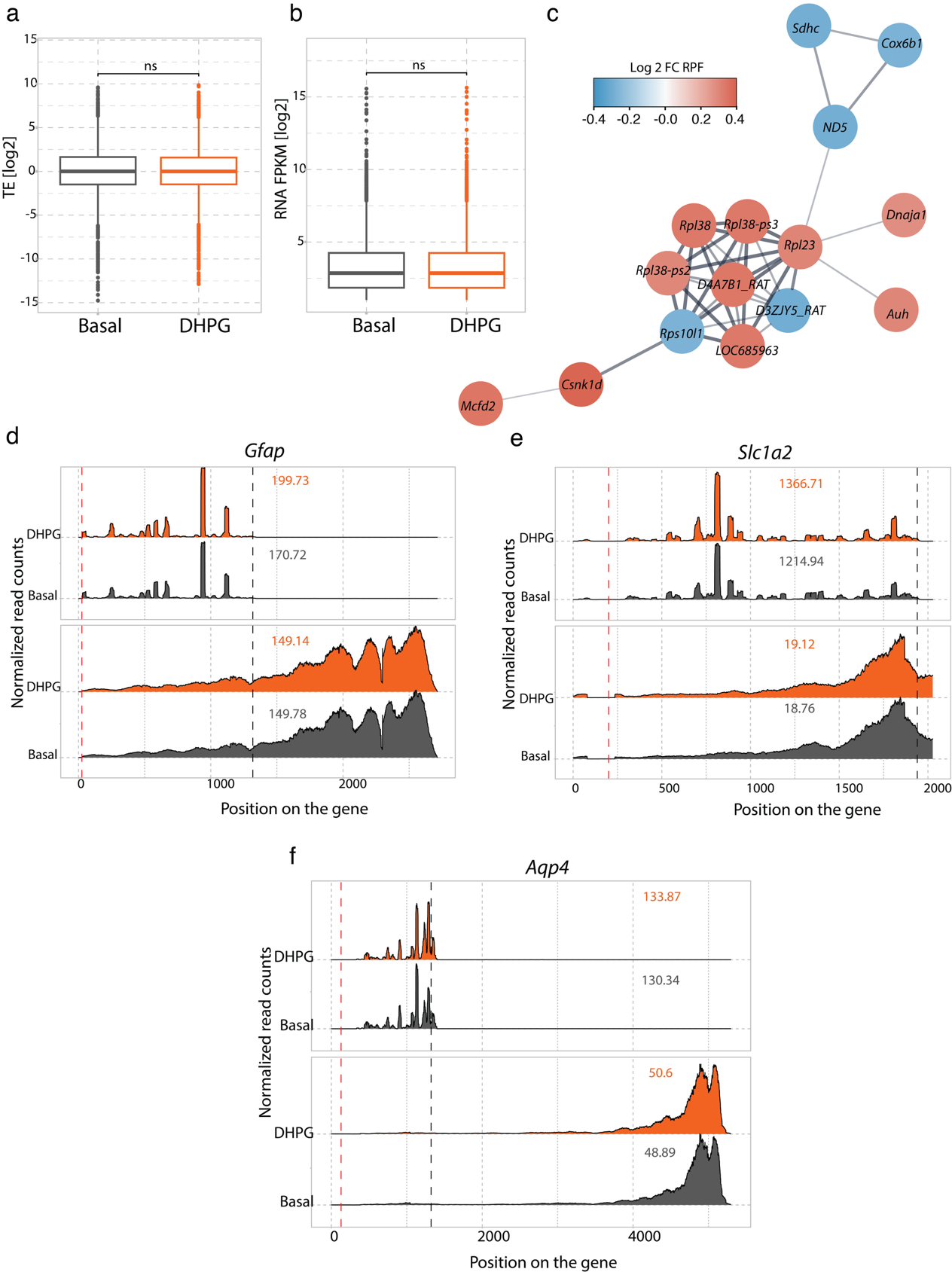

**Supplementary figure 2 (related to figure 2): Differential gene analysis at the TE and RNA level, cytoscape**

a. Log2 of TE  between basal and DHPG treated samples
a. Log2 of RNA (FPKM) between basal and DHPG treated samples
a. Cytoscape analysis of differentially expressed transcripts gave a major cluster of ribosomal and mitochondrial transcripts.
a. Gene silhouettes for Glial fibrillary acidic protein (*Gfap)*, marker for Glia, for RPFs and total RNA libraries. Red and black dotted lines indicate translation start and termination sites respectively.
a. Gene silhouettes for Solute carrier family 1 member 2 (*Slc1a2*), glial marker, for RPFs and total RNA libraries. Red and black dotted lines indicate translation start and termination sites respectively.
a. Gene silhouettes for Aquaporin 4 (*Aqp4*), an glial marker, for RPFs and total RNA libraries. Red and black dotted lines indicate translation start and termination sites respectively.

**Supplementary Table 1: List of differentially regulated transcripts at the RPF level**

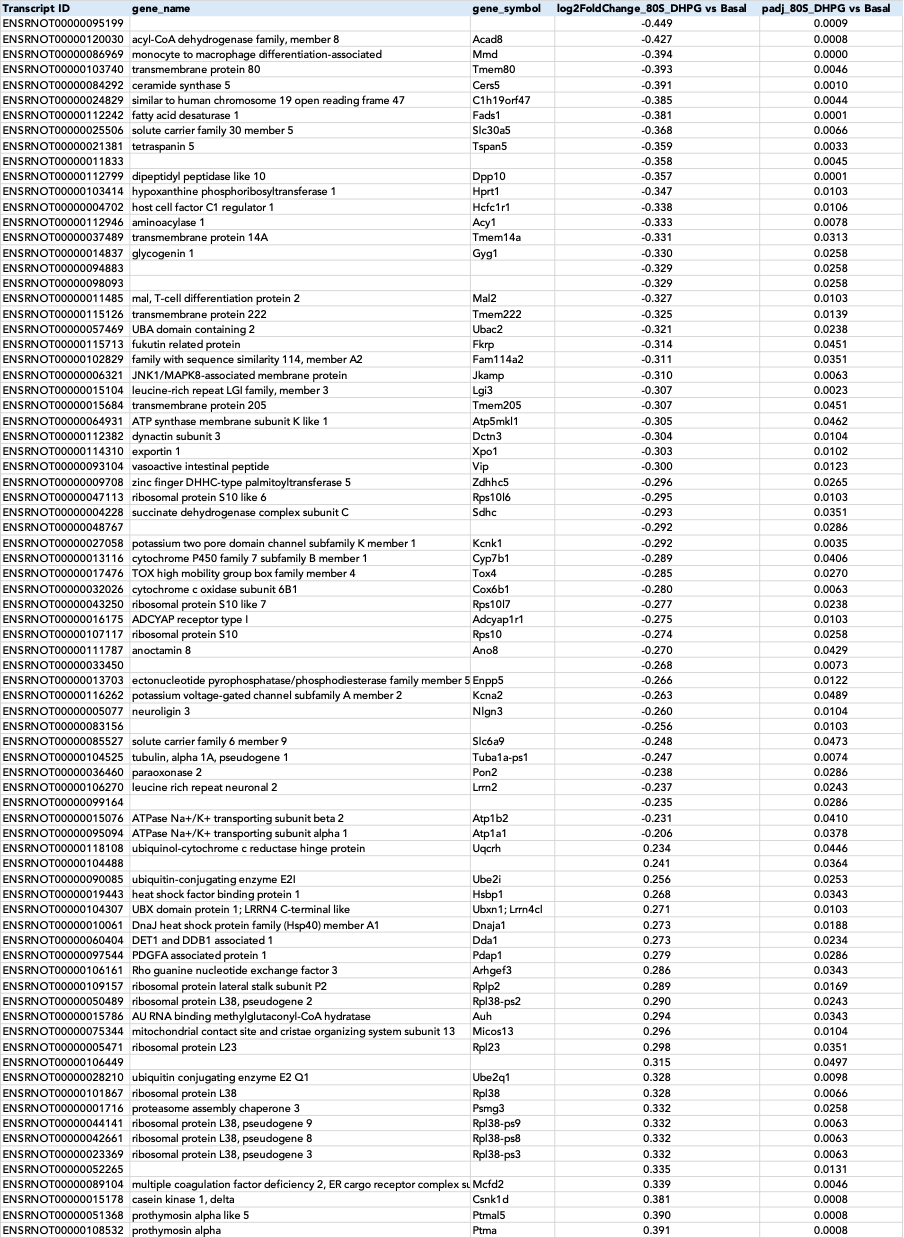
